# Compositional Differential Abundance Testing: Defining and Finding a New Type of Health-Microbiome Associations

**DOI:** 10.1101/2024.06.04.596112

**Authors:** Siyuan Ma, Curtis Huttenhower, Lucas Janson

## Abstract

A major task of microbiome epidemiology is association analysis, where the goal is to identify microbial features related to host health. This is commonly performed by differential abundance (DA) analysis, which, by design, examines each microbe as isolated from the rest of the microbiome. This does not properly account for the microbiome’s compositional nature or microbe-microbe ecological interactions, and can lead to confounded findings, i.e., microbes that only appear to associate with health through their confounding association with health-related, biologically informative microbes. To remedy these issues, we present Compositional Differential Abundance (CompDA) analysis, a novel approach for health-microbiome association. CompDA provides a novel approach to identify health-related microbes by examining the microbiome holistically, which a) accounts for the data’s compositionality and ecological interactions, and b) has clear interpretations corresponding to host health as affected by microbiome-based interventions. CompDA prioritizes health-related microbes and controls false discoveries by implementing recent advances from high-dimensional statistics, and can be flexibly adapted to many common tasks in modern microbiome epidemiology, including enhancing microbiome-based machine learning by providing rigorous p-values to prioritize important features. We validate the performance of CompDA, and compare against canonical microbiome association methods including DA with extensive, real-data-informed simulation studies. Lastly, we report novel and consistent findings of CompDA in application, based on re-examination of recently reported microbial signatures of colorectal cancer in a meta-analysis.

## Introduction

Population-scale studies of the human microbiome aim to identify microbial elements that are important to host health^1,2^, with findings associating dysbiotic microbiomes with various diseases such as diabetes, inflammatory bowel diseases, and cancer^3–5^. A key component of these efforts is microbiome-appropriate statistical analysis^6^. This includes, importantly, differential abundance (DA) analysis^7^, which tests for microbial features (taxa, gene families, pathways) that are individually associated with host health. Notably, however, none of these widely used methods account for any type of non-independence between microbes. This can include the ubiquitous compositionality of nearly all microbiome measures, as well as ecological interactions such as metabolic dependence. Ignoring such dependencies results in DA biomarkers that include truly health-associated microbes as well as spurious findings^8–10^ (covarying only due to compositionality or other types of covariation).

A number of microbiome-specific and general ‘omic challenges arise when analyzing microbiome data, including (1) confounding microbe-microbe associations, (2) compositionality, and (3) high-dimensionality. First, microbial feature abundances are associated with each other from ecological interactions^12^. This contributes to spurious health associations, because while some microbes are truly related to health, there are “spurious” microbes that do not interact with health but correlate with the health-related ones. The second group will have confounded, false associations with health^13^, similar to non-causal linkage loci in genome-wide association studies^14^. Second, in most application settings, microbial abundances are measured on the relative or “compositional” scale^11^. This also leads to issues of true versus false microbes associated with health, as the increased abundance of a microbe under a certain condition necessarily implies decreased abundance of some other microbe(s). Third, microbiome studies, same as other molecular epidemiology applications, are high-dimensional (number of features in the 100s to 1000s, greater than number of subjects in the 10s to 100s). Advanced approaches such machine learning are often desirable to achieve good representation of the health-microbiome relationship in this setting, but it is challenging to evaluate the statistical significance of microbes deemed as “important” by these machine learning models, thus posing challenges for controlling false discoveries^15^. Statistical methods must account for these spuriousness mechanisms to generate robust findings for the microbiome.

Apropos, most commonly adopted DA frameworks examine microbe-health associations on an individual-microbe basis, and do not account for the rest of the microbiome^16^. This makes it difficult to differentiate between health-associated, biologically informative microbes versus covariation relationships, either due to confounding microbe-microbe association (a non-health related microbe appears to have association with health, through its association with microbes directly impacting health), or the compositional effect of microbiome data (an increase in the abundance of health-related microbes will necessarily lead to decrease in non-health-related abundance), i.e., the first two challenges highlighted above. Recent developments in the microbiome field aim to address the compositionality issue for DA analysis (e.g., LinDA^17^ and robust DA^18^), but still operate within the individual microbe framework. In practice, DA approaches are most appropriate for providing initial candidate hits for further investigation. Recent research in the field suggests the need for more judicious alternatives^8–10^. We suggest adopting a holistic, multivariable approach, which evaluates health-microbe associations while accounting for the entire microbiome and its dependency structures to eliminate spurious findings. Such methods have been developed for human genetic studies, to fine-map true causal loci of phenotypic traits from potential spurious associations due to linkage, and have become an integral step for refining initial hits in genome-wide association studies^19,20^. Similar efforts for microbiome data are limited^21–24^, likely due to a fundamental, data-unique challenge. When testing each microbe, these methods directly adjust for all other microbes, and constrain the microbe of interest’s relative abundance to a fixed constant (one minus the abundance of others). Any potential association between a microbe versus host health is thereby nullified, because a fixed constant by definition does not have associations with any outcome. The aforementioned multivariable methods for microbiome data do not address this issue, impacting their interpretability. For adopting multivariable testing to control false discoveries in microbiome research, advancement is required for a) conceptually well-defined microbe-health associations that are meaningful for compositional microbial relative abundances, and b) in practice, a modeling framework and implementation to test for such associations while controlling false discoveries.

In this work, we propose a novel multivariable testing framework that reduces spurious findings and prioritizes biologically informative health-microbe associations. We first provide a new multivariable definition for *compositional* microbiome-health associations that is meaningful for microbial relative abundances and has a clear clinical interpretation. We then generalize a novel framework for multivariable testing, namely, the conditional randomization test (CRT)^15^, for a practical implementation while better accounting for data-specific characteristics of the microbiome (zero-inflation, compositionality, ecological dependencies). With real-data-informed simulation studies, we validate our method and compare against existing microbiome testing methods and establish its performance in achieving power while controlling false discoveries. We also apply our work to a real-world study to identify biologically informative microbial associations with colorectal cancer^25^ and deprioritize spurious microbes. We provide a publicly available implementation of our method, named CompDA (Compositional Differential Abundance testing). To our knowledge, our work represents one of the first *multivariable* testing efforts appropriate for microbiome data, thus achieving good control of confounded, “false discovery microbe” findings. In practice, it provides the theoretical framework, methodology, and implementation for prioritizing true biomarkers for microbiome translational research.

## Results

### CompDA: compositional microbiome multivariable testing to prioritize health-related, true-positive microbes

We propose CompDA, a holistic (i.e., multivariable) and compositional testing framework as an alternative to traditional DA testing, to better eliminate spurious findings and prioritize the most biologically informative microbial markers (due to their conditional dependence, **Figure 1**). It makes three important contributions. First, CompDA provides a novel, clinically meaningful definition for health-related microbes by adapting traditional multivariable testing to both adjust for confounding microbe-microbe associations and account for the data’s compositionality. We propose to modify existing multivariable methods with a novel definition for *compositional* health-microbe association, where each microbe is considered as health-related when associations exist while adjusting for the “balance” of the other microbes (i.e., renormalized relative abundances, **Figure 1B**). Mathematically, our approach performs modified multivariable statistical tests, to both accommodate the microbiome’s compositionality constraint (because each microbe can freely vary while holding the balance of other microbes constant), and control for confounding microbe-microbe associations (**Methods**). More importantly, for biological and clinical interpretations, our novelly defined compositional health-microbe associations directly correspond to host-health-related interventions on pathogenic or commensal microbes. Introducing commensal microbes (e.g. through fecal microbiome transplantation, or FMT^26^) has been studied as an emerging treatment for certain health conditions. In the simplest case, to study the effect of an individual microbe towards a health outcome, a randomized trial will introduce the microbe to subjects in the treatment arm. This corresponds to modifying the abundance while holding the renormalized “balance” of other microbes constant, because introducing a microbe to an environment also changes the other microbes’ relative abundances but not their balance. The argument generalizes easily beyond single microbes to cocktails of commensals as interventions. Thus, true health effects in this microbial intervention regime correspond exactly to CompDA’s definition for compositionally health-related microbes, highlighting its interpretation in meaningful clinical context.

**Figure 1:**
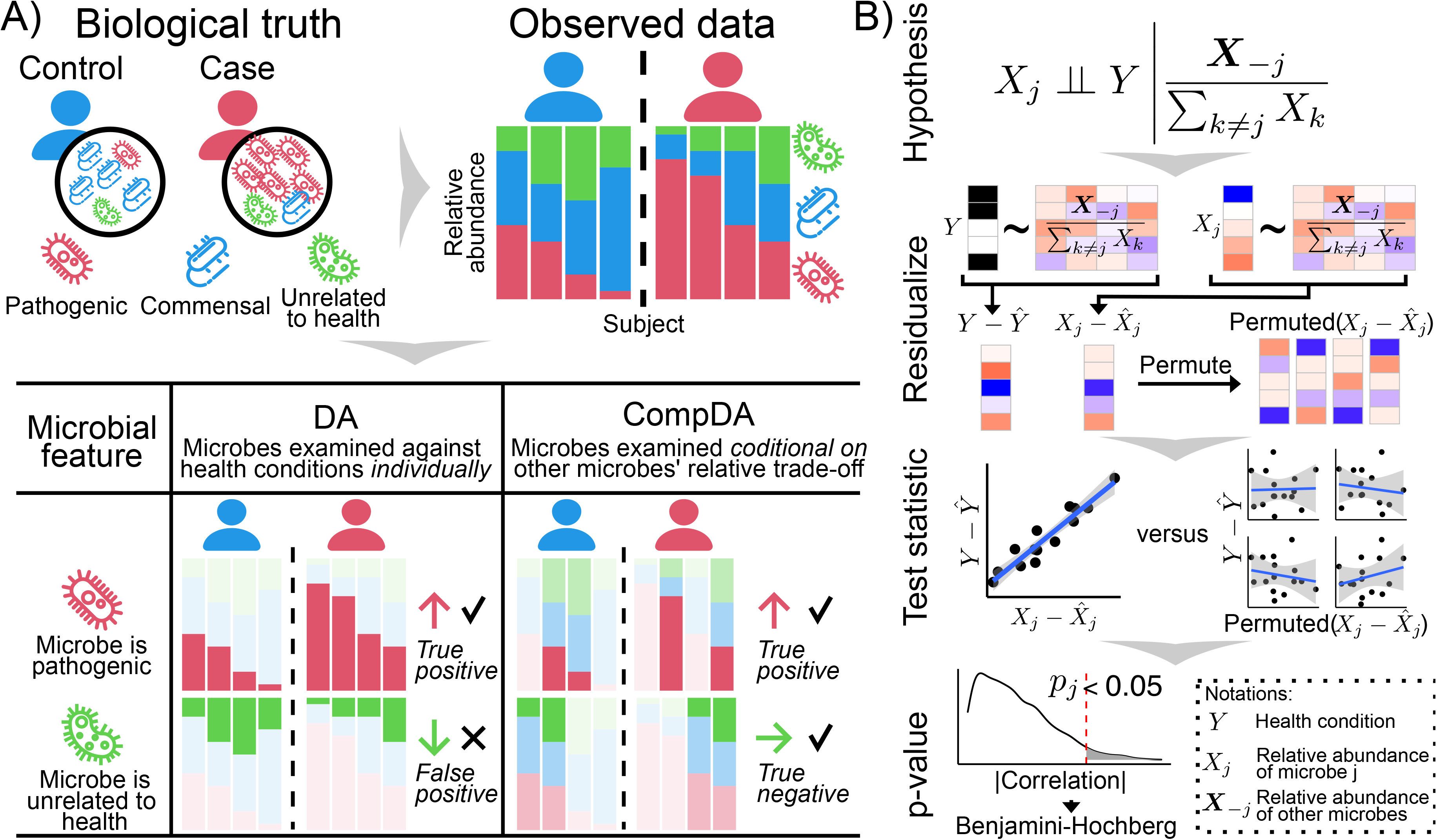
CompDA prioritizes true, health-related microbial markers and controls false discoveries. **A**) Traditional DA analysis does not differentiate between true, health-associated microbes versus spurious findings. In the illustrated example, the pathogenic microbe (red) is enriched in disease conditions and the commensal microbe (blue) is depleted. However, due to the data’s compositional nature, a microbe unrelated to health (green) also displays differential relative abundances with respect to health, and will be reported as a false positive finding by DA analysis. In comparison, CompDA accounts for the rest of the microbiome when testing each microbial feature, thus correctly prioritizing true health-related features and eliminating spurious findings. **B)** Overview of CompDA’s novel definition of compositionally health-related microbes, and corresponding methods to identify them. First, CompDA defines a new type of statistical null, whereby a microbe is only considered associated with health while conditional on the renormalized abundance of other microbes. This appropriately accounts for confounding microbe-microbe associations and the data’s compositional nature (**Results, Methods**). To test for this new type of compositional association across many microbes (i.e., high-dimensional data) while controlling false discovery rate, CompDA then conducts conditional randomization testing^27^ for each feature. Both the health outcome and the microbe under investigation are residualized against the renormalized abundances of other microbes, thereby eliminating potential confounding associations explainable by the other microbes. The correlation between the residuals is used as the test statistic to contrast against its permuted null distribution, which provides valid p-values that correctly control false positives. Lastly, the p-values across microbes are synthesized with false discovery controlling procedures such as Benjamini-Hochberg, which correctly prioritizes true health-related microbes and eliminates false discovery findings.

Second, we achieve control of false discoveries among many candidate microbial features by adopting recent advances in high-dimensional statistics research^27^ (**Figure 1B, Methods**). The typical dimensionality of microbiome studies (similar or greater number of microbial features than sample size) poses the “high-dimensional” challenge in statistical inference, where multivariable testing has seen promising advances in recent years in robust ways to control false discoveries^15^. For domain-specific health-microbiome multivariable testing, we build upon the recently proposed conditional randomization testing (CRT^27^) approach. Intuitively, the CRT contrasts observed microbe-health associations (accounting for other microbes) versus artificial, randomized “null” associations (again, accounting for other microbes), and thus provides valid p-values for statistical inference (**Methods**). Our contribution is to tailor the randomization procedure towards microbiome data, accounting for its compositional, zero-inflation, and ecological interaction natures (**Methods**). This ensures valid, false-positive-controlled p-values for each microbe, which are further synthesized with false discovery rate (FDR) control procedures to prioritize true health-associated microbes among hundreds to thousands of candidates and control false discoveries.

Lastly, CompDA is flexible in various practical settings. First, it can handle different types of health-related outcome variables (binary, continuous, count-based). This is because CompDA is consistently valid for different types of outcome variables. In particular, it can adopt various test statistics appropriate for the particular type of outcome under investigation (**Methods**). Second, confounding covariates can be easily included in CompDA testing in addition to the balance of other microbes (**Methods**), similar to traditional DA analysis. Third, CompDA can be combined with different microbiome-based prediction models, including state-of-the-art machine learning^28,29^, to provide valid p-values as principled measures of microbe importance. In summary, CompDA is a comprehensive and flexible testing framework for microbiome epidemiology, that eliminates spurious finding and prioritizes biologically informative microbe-health associations.

### CompDA outperforms both DA analysis and existing multivariable methods to prioritize health-related microbes

In extensive, real-data-based validations, CompDA successfully eliminated spurious microbes to prioritize true, health-related targets, when compared against traditional DA testing as well as previous multivariable testing paradigms designed for microbiome data (**Figure 2, Supplemental Figures 1-2**). Our simulation was strongly informed by real-world studies, where the microbiome data were directly obtained from publicly available studies (as opposed to generated *in silico*). Only the relationship between the health outcome and the microbiome was simulated (**Methods**). This ensured that our conclusions were drawn upon (and thus generalizable to) real-world microbial communities, especially regarding confounding microbe-microbe associations. Across our simulation studies, we observe that CompDA consistently outperformed both DA and high-dimensional microbiome testing methods, and successfully controls false discovery rates while achieving good power.

**Figure 2:**
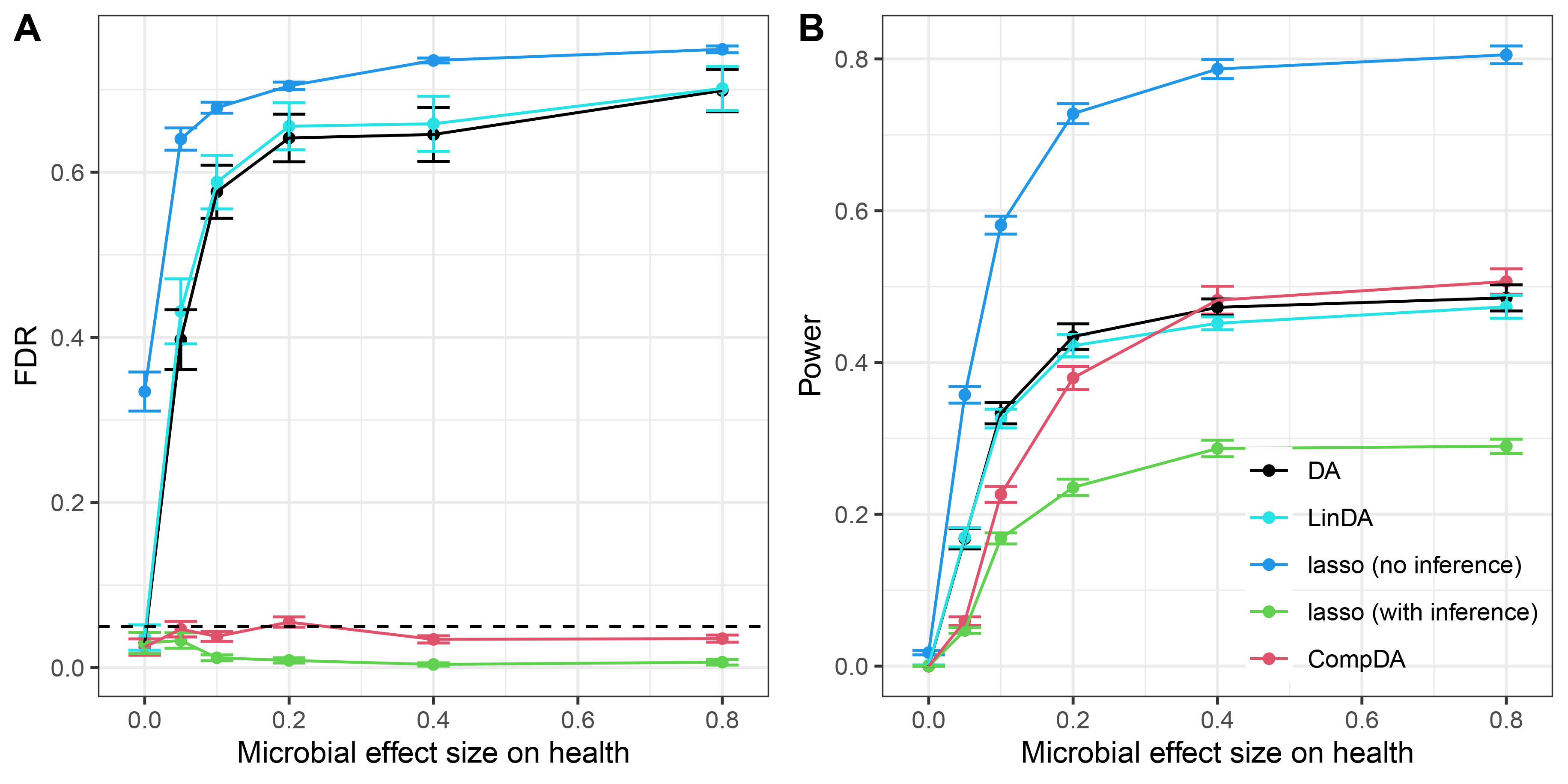
CompDA achieves valid and powerful high-dimensional testing of multivariable, compositional health-microbiome associations. We evaluated the performance metrics of different methods including false discovery rates (**A**) and power (**B**), varying different simulation parameters while holding others constant. The presented results in the figure evaluate various health-microbiome association effect sizes while fixing sample size (400), complexity of community (200 microbes), and signal density (10% microbes associated with health). Results varying the other simulation parameters are presented in **Supplemental Figures 1-2**. Across various strengths of health-microbiome association effect sizes, CompDA consistently controls the false discovery rate below the targeted threshold of 0.05, whereas traditional DA analysis, compositionality-adjusted DA (LinDA) and lasso (selecting non-zero coefficients without the appropriate statistical inference) suffers severe FDR inflation at higher effect sizes. Among the methods that are able to control the FDR, CompDA also achieves the best power. It is thus the only method that successfully controls false discovery findings and prioritizes true, health-related microbes.

In the representative example of identifying microbes associated with case/control binary health conditions, CompDA successfully controlled false discovery rates, and achieves the best power compared to other methods that were also able to do so (**Figure 2**). Starting from 8,000 healthy, western stool samples obtained from publicly available data^30^, we simulated disease-control binary outcomes associated with random subsets of the microbiome (**Methods**). We varied a wide range of simulation parameters, including the true effect size (magnitude of microbial effects on health), signal density (the number of true health-related microbes), and data dimensionality (both sample size and the number of microbes). CompDA successfully controlled the false discovery rate under the targeted threshold of 0.05. This was in contrast to existing DA testing^7^, compositionality-adjusted DA testing (LinDA^17^), or high-dimensional variable selection through the ubiquitous lasso regression^31^, all of which demonstrated severe FDR control issues especially with large health-microbiome association effects. The existing method of testing for coefficients from lasso regression (lasso with inference) was also able to control the false discovery rate but has substantially reduced power. We emphasize that DA paradigms (including compositionality-adjusted methods such as LinDA) cannot handle such tests well by design, because they cannot differentiate between biologically informative, health-related microbes, versus those that are only spuriously associated with health because they are also associated with the biologically informative microbes. This highlights how CompDA’s novelly defined compositional health-microbe association is specialized for microbial relative abundances. CompDA’s performance is consistent across simulation scenarios, including different types of health outcomes, health-microbe signal densities, sample sizes, and microbial community complexities (**Supplemental Figures 1-2**). In practice, a CompDA analysis can be conducted at around ten minutes on a typically sized microbiome dataset with single-core computation (**Supplemental Figure 3**), with parallelization available in our implementation for additional computation speed. We thus conclude that, in common health-microbe association applications, CompDA was the only method that successfully controlled false discoveries, and prioritized health-related, biologically informative microbes with high power.

**Figure 3:**
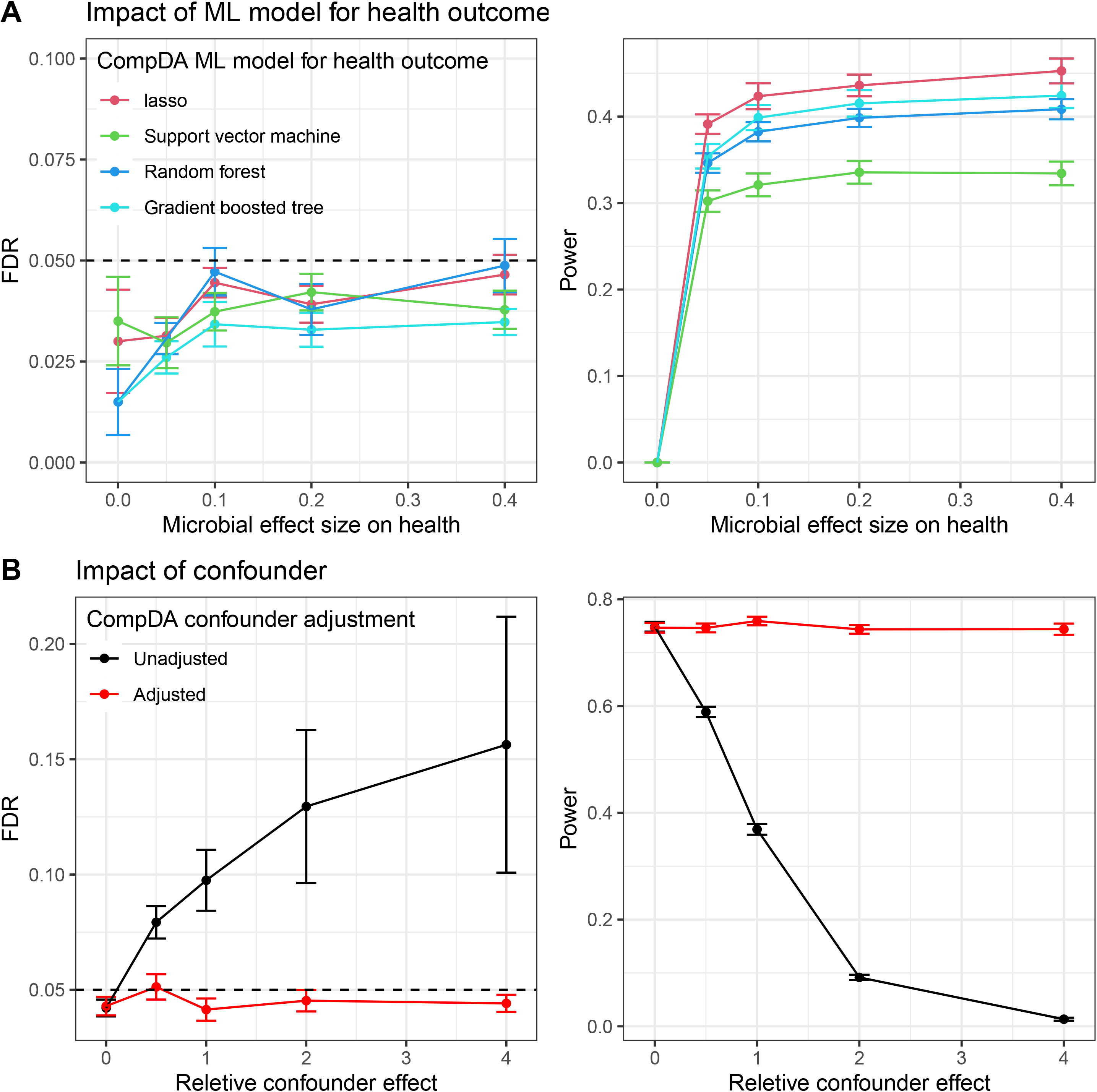
CompDA performs microbiome machine learning with robust performance and can flexibly adjust for host confounders. **A**. CompDA provides valid p-values for features selected from microbiome machine learning paradigms. This performance is robust when adopting different machine learning modeling choices for modelling the health outcome given the microbiome, including lasso regression, support vector machine, random forest, and gradient boosted tree. This holds true even when such models are misspecified, i.e., they disagree with the true data generation mechanism. **B**. CompDA controls the false discovery rate and maintains power when health-microbiome relationships are confounded by host factors (age group). This is realized by accounting for such host factors in CompDA modeling, just as with regular DA analysis. The false discovery rate is consistently controlled at varying levels of confounding strength, as long as such confounders are accounted for in the modeling.

### CompDA enhances microbiome machine learning and can flexibly adjust for confounders

CompDA has a flexible implementation that is applicable to various microbiome epidemiology tasks with robust performance (**Figure 3**). In recent years, microbiome studies increasingly adopt modern machine learning to identify microbial signature patterns associated with host health^25,28,29^. CompDA can flexibly incorporate such advances by swapping in any health-microbiome prediction model of choice (**Methods**) while filling two important gaps (**Figure 3A**). First, while existing approaches output important microbial features from machine learning, they do not provide confidence assessment on the selected features, e.g., statistical p-values with guarantees to control false positives. Consequently, microbial signatures reported to be health-associated by these methods are often prioritized by some arbitrary threshold, which can severely violate false discovery controls. For example, random-forest-based methods report the top X microbes ranked by variable importance^29^, but there is no theoretical guidance on how X should be determined. With an ill-selected threshold, the prioritized microbes can be severely inflated for false discoveries. As we clarify in **Methods** and validate with simulations (**Figure 3A**), CompDA provides valid statistical significance when adopting modern machine learning in modeling health-microbiome relationships; it can thus confidently and flexibly prioritize health-associated microbes as predicted by machine-learning models, while controlling false positive findings. Second, it is unrealistic for any machine learning model to perfectly capture the relationship between host health and the microbiome. In this regard, CompDA is also robust against model misspecification. That is, CompDA’s p-values for health-microbiome relationships are valid when adopting different machine learning models, even when they deviate from the biological truth. We again validate this with simulations, where the machine learning models selected for CompDA testing were intentionally set to be different from the true data generating mechanism (**Methods**), but our method still consistently controlled false discoveries (**Figure 3A**). Overall, CompDA enhances microbiome machine learning by performing valid, robust, and flexible prioritization of health-related microbes while incorporating modern machine learning models.

Microbiome epidemiology often requires adjustment for potentially confounding host and environmental factors, for example in regular DA analysis^7^. CompDA can robustly achieve this just as state-of-the-art DA methods, which we validate with simulation studies (**Figure 3B**). We included subject age, known as associated with both the microbiome and host health, thus constituting a confounder for health-microbiome associations^32^. As in our primary simulation studies, the age variable used here is directly obtained from real-world data (along with the microbiome and their inter-relationships thereof, **Methods**); only the effect of host factors on health is simulated. This strengthens the generalizability of our simulation results towards real-world applications. As presented in **Figure 3B**, CompDA successfully accounts for confounding the host factor and controls the false discovery rate, while maintaining statistical power to identify true, confounder-adjusted microbes related to health.

### CompDA identifies meaningful microbial signatures for colorectal cancer

We applied CompDA to study microbial signatures of colorectal cancers (CRC), where the field made recent advances towards microbial biomarkers of the disease^25,33^. By employing a collection of publicly available metagenomic sequencing studies of CRC^25^ (**Methods**), we applied CompDA to identify disease-associated microbial species, and compared our findings with that of DA analysis (**Figure 4**). CompDA selected three species associated with CRC at an FDR threshold of 0.05: *Fusobacterium nucleatum, Peptostreptococcus stomatis*, and *Streptococcus salivarius* (**Figure 4A**). These were all identified as important, predictive microbes for CRC through machine learning in the original publication^25^. CompDA’s main contribution here is to enhance microbiome machine learning by providing rigorous, statistical p-values on the important predictive features, such that researchers can confidently prioritize health-associated microbes with non-arbitrary thresholds that guarantee, e.g., controlled false discovery rates.

**Figure 4:**
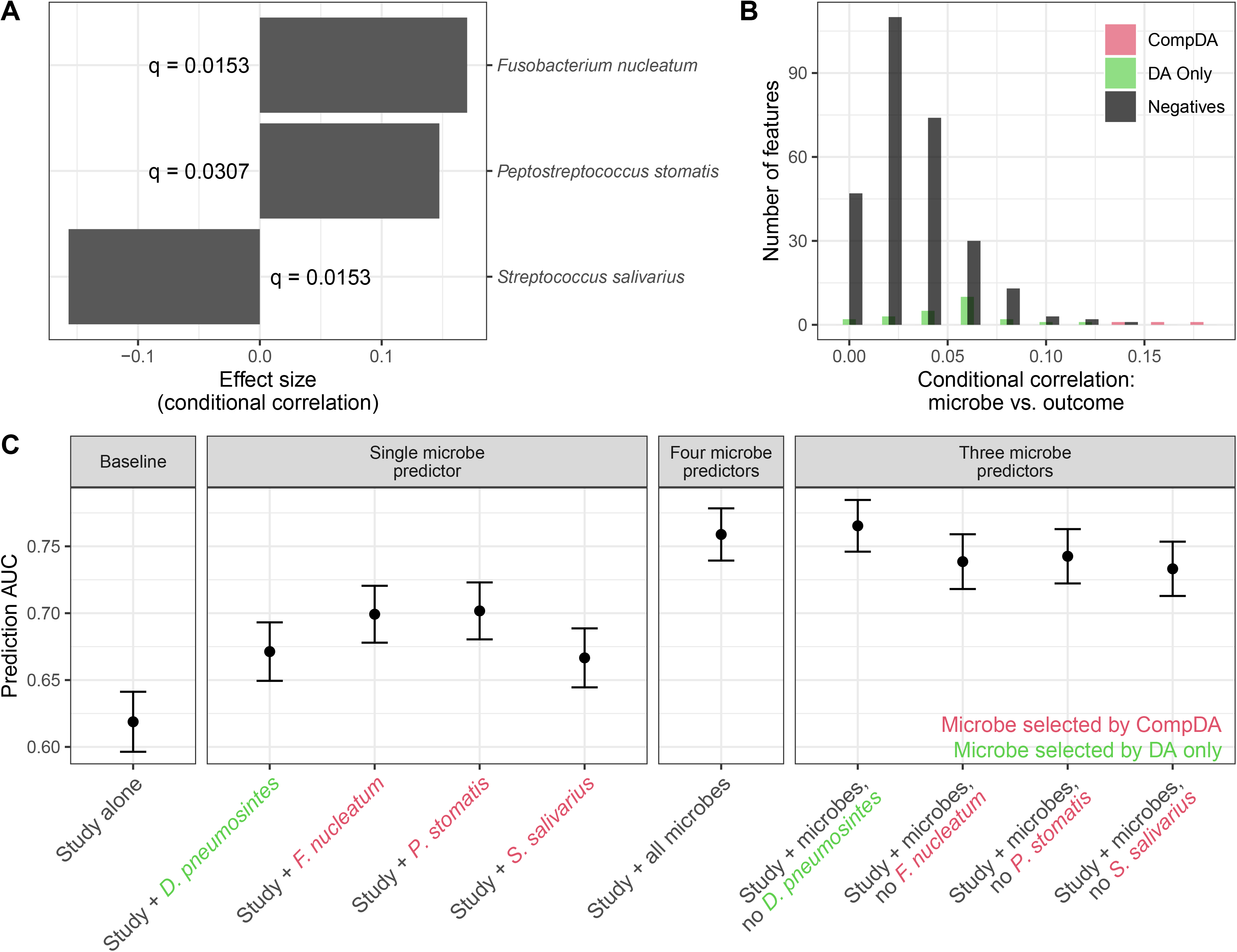
CompDA prioritizes microbes associated with colorectal cancer and eliminates spurious findings from DA analysis. **A**. CompDA identified three microbial species confidently associated with CRC (FDR-adjusted q < 0.05). All three microbes were reported as important signatures via microbiome-based machine learning; CompDA establishes rigorous guidance to prioritize such features by providing statistical p/q-values. X-axis: conditional correlation of each microbe associated with CRC status when accounting for the rest of the microbiome (**Methods**). **B**,**C**. CompDA eliminates spurious, false positive microbes reported by DA analysis. Microbes selected to be associated with CRC (q < 0.05) only by DA had much smaller (absolute) conditional correlation with the disease compared to those identified by both CompDA and DA (**B**). This suggests that microbes selected only by DA contain many spurious associations with CRC, which would be explained by accounting for the rest of the microbiome (thus including the true CRC-related microbes). We provide such an example in *D. pneumosintes*, a microbe reported by DA that is likely spurious (**C**). While *D. pneumosintes* is associated with CRC by itself, this association is largely explained by the CompDA-selected microbes (*F. nucleatum, P. stomatis, and S. salivarius*). As shown by the predictive area under the curve (AUC), each microbe is individually predictive for CRC. However, a prediction model for CRC status with all four microbes has similar performance versus if only the three CompDA-selected microbes were included, whereas each microbe selected by CompDA still contributes to predictive performance even the other microbes were included. This suggests that CRC-related signal in *D. pneumosintes* is likely spurious and can be explained by the CompDA-selected microbes.

Compared against traditional DA analysis, CompDA effectively prioritizes biologically informative, CRC-related microbes while reducing spurious associations (**Figure 4B,C**). The conditional associations of microbes with CRC (conditional upon the rest of the microbiome, see **Methods** for details) is markedly higher in features prioritized by CompDA compared to DA (selected with q < 0.05 by both, **Figure 4B**). This suggests that many microbes identified by DA analysis are correlated with other microbes that more directly related to CRC. We highlight such an example in the case of *Dialister pneumosintes*, a spurious microbe given the true biomarkers (*F. nucleatum, P. stomatis, and S. salivarius*; **Figure 4C**). *D. pneumosintes* is identified as highly differentially abundant with respect to CRC by DA analysis (FDR adjusted q = 8.12 x 10^-9^) but not by CompDA (q = 0.953), suggesting that it has spurious associations with CRC. Across *D. pneumosintes* and the three microbes selected by CompDA, we performed predictive modeling of CRC using various subsets or full set of the four microbes, additionally including study indicator as a baseline predictor. Individually, all four microbes had predictive power for CRC (panel “single microbe predictor”). However, once the three CompDA-selected microbes were included in the model, *D. pneumosintes* does not provide additional prediction power (panels “four/three microbe predictors”), suggesting that DA-identified *D. pneumosintes* is likely a spurious association, accounted for by other microbes. This confirms that *F. nucleatum, P. stomatis, and S. salivarius* are likely true microbial markers for CRC, whereas *D. pneumosintes* is likely only spuriously associated with CRC only through the other microbes. CompDA thus uniquely separates true, CRC-associated microbial signatures from spurious, false microbes, compared to traditional DA analysis.

## Discussion

Here we propose CompDA, a novel method for testing health-microbiome associations, that accounts for the data’s compositionality, confounding microbe-microbe associations, and high-dimensionality. As an improvement to traditional DA analysis, CompDA aims to eliminate microbes that can be erroneously reported as health-related through their confounding associations with true-health related ones. Thus, the microbes prioritized by CompDA are well controlled for false discovery rates; for interpretation, they also correspond to microbial treatment effects in trials of such interventions. We validated CompDA’s performance with extensive real-data-based simulation analysis and showcased its utility with a real-world application. CompDA thus provides a more judicious alternative to traditional DA analysis in testing health-microbiome relationships, that has a novel (and in certain settings, improved) clinical interpretation.

A desirable feature of CompDA is that it is robustly valid (i.e., control of false discoveries) even when the relationship between host health and the microbiome is misspecified. It can thus be flexibility applied to various application settings, including in the context of microbiome-based machine learning, without concerns of whether the postulated machine learning model deviates from the biological truth. Indeed, the method’s validity is empirically robust against model misspecification (**Figure 3A**). This is because our conditional randomization testing procedure ensures that CompDA can control the false discovery rate even when health-microbiome relationships are misspecified. As a consequence, for modern prediction modeling of the relationship between host health and the microbiome (e.g. with random forest or support vector machine^28^), CompDA can consistently provide valid statistical tests for the importance of individual microbial features, regardless of the adopted model. Such efforts are gaining popularity in recent years^29^, but lack a principled way to prioritize the important microbes identified from model training. CompDA enhances existing microbiome machine learning efforts by providing a statistically justified way to identify important microbes that controls false discoveries and remains consistently valid for different models of choice.

Computation cost is an important consideration for high-dimensional data modeling problems including those of the microbiome. To ensure the scalability of CompDA, we employed recent advances in high-dimensional conditional testing^27^, such that only one iteration of high-dimensional model fitting is required for testing each microbe. In practice, we observe that, for regularly sized microbiome datasets, data fitting can be achieved within reasonable time even with single core computing (**Supplemental Figure 3**). This can be further improved when multiple computation cores are available, as the CompDA procedure is fully parallelizable and is implemented as such.

There are several aspects of CompDA that call for extensions. First, although CompDA’s validity consistently holds under different relationships between health and the microbiome, it does require that the microbiome observations are independent between each other. This limits our application to cross-sectional designs. Under e.g. longitudinal settings, because different timepoints’ microbiomes are presumably correlated with each other, CompDA’s validity no longer holds. Second, it is difficult to test for interaction effects between pairs of microbes jointly on health. Interaction testing is a notably difficult problem for molecular profiles (e.g., gene-environment interaction testing^34^). For future work, it is of interest to develop specialized procedures towards such interaction effects.

## Methods

### Novel multivariable testing to prioritize true health-related microbes

CompDA proposes a novel multivariable test to identify health-related microbes, while eliminating spurious findings and accounting for the data’s compositional nature. Let *x* = (*x*_1_,*x*_2,…,_*x*_*p*_) denote the *p*-dimensional vector of microbial relative abundances, and *Y* denote a continuous (e.g., BMI) or binary (e.g., disease status) health condition of interest. In the simple case of no confounding host or environmental variables (which is without loss of generality - we demonstrate later that our theory and methodology easily incorporates the presence of covariates), existing differential abundance testing methods examine the marginal hypotheses *Y X*_*j*_,That is, health is not marginally associated with the *j*-th microbe. While useful in providing an initial list of candidate microbial markers for health, this approach does not control false positives. This is because for both health-related microbes and spurious microbes that interact with the health-related ones but not health itself, this null does not hold (**Figure 1A**). As such, current differential abundance testing methods cannot distinguish between true, health-related microbes versus spurious ones, and generate false discoveries.

Alternatively, existing multivariable modeling, such as multivariable regression, is not directly appropriate for microbiome data. To illustrate this, classical multivariable regression examines the association between and with a (generalized) linear model E*Y* = *g* (*β*_1_*X*_1_+ *β*_2_*X*_2+_…*+ β*_1_*β*_*p*_) Here *β*_1_ ,…, *β*_*p*_ are regression coefficients for each microbial feature and *g* is the link function. For a microbe *X*_*j*_, a zero-regression coefficient *β*_*j*_ = 0 indicates conditional independence: *X*_*j*_ *Y*| *X*_*−j*_, where *X*_*−j*_ indicates the vector of microbial abundances excluding feature *j*. For regular data types, this indicates that the *j*-th feature is not related to the health condition while adjusting for others. For microbiome data, the compositionality constraint *X*_*j*_ = 1−∑*X*_*−j*_ applies. Here we adopt a slight misuse of notation and use ∑*X*_*−j*_ to denote ∑_*k*≠*j*_ *X*_*k*_ for simplicity. Thus the conditional null *X*_*j*_ *Y*| *X*_*−j*_ always trivially holds, because *X*_*j*_ is a *constant* conditional on *X*_*−j*_ and independent of any random variable. This means that existing multivariable testing methods, traditionally used for removing spurious findings, are not meaningful for the microbiome.

We propose a novel conditional null for multivariable testing of health-microbiome associations. Specifically, for an individual feature *X*_*j*_, we propose to examine the following conditional null:

#### **Definition 1**.

A microbe is conditionally not associated with health if its relative abundance is independent with the outcome while conditioning on the renormalized abundance of the rest of the microbiome. That is,

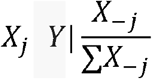

Here the “balance” of the rest of the microbiome is defined through renormalization of the conditional set, i.e.,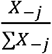. Our approach is thus uniquely tailored towards compositional microbiome data, and investigates candidate health-related microbes, by accounting for the balance of the rest of the microbiome (**Figure. 1**).

**Definition 1** defines a meaningful set of health-related microbes, i.e.,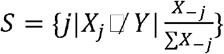 (the set of microbes where the conditional null in **Definition 1** does not hold). This definition makes three novel contributions. First, mathematically, it precludes spurious false discoveries by conditioning on the rest of the microbiome when examining health-microbe associations. Spurious microbes that are not directly associated with health, but interact with true, health-related ones are conditionally independent of health, and will not be included as potential microbial markers. This is similar to traditional confounder adjustment in, e.g., multivariable regression, and GWAS fine mapping for identifying true causal loci^14^.

Second, from a technical standpoint, **Definition 1** bypasses the compositional constraint of microbial relative abundances, as each microbe under investigation can still freely vary under the conditioning operation. True, health related microbes will not be trivially deemed unimportant for any outcome, unlike under the hypothesis tested by existing multivariable approaches.

Last, **Definition 1** importantly characterizes health-microbiome associations that are biologically and clinically meaningful, as the defined relationships correspond directly to health-related microbial intervention. Introducing commensal microbes (e.g. through fecal microbiome transplant, or FMT^26^) has been studied as emerging treatments for certain health conditions. In the simplest case, to study the causality of an individual microbe *X*_*j*_ towards a health outcome *Y*, a randomized trial will introduce the microbe to subjects in the treatment arm. This corresponds mathematically to modifying the abundance *X*_*j*_ while holding the renormalized “balance” of other microbes 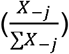 constant, because introducing a microbe to an environment also changes the other microbes’ relative abundances but not their balance. Thus, a null causal effect in this microbial intervention trial corresponds exactly to 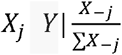 .(**Definition 1**). This establishes the equivalence between our proposed novel conditional health-microbe associations and clinically meaningful microbial causality, defined through the above intervention regime. The argument generalizes easily beyond single microbes to cocktails of commensals as interventions; our novel definition thus corresponds to meaningful clinical context, e.g., causal microbial effects in FMT trials.

### Conditional randomization testing of multivariable health-microbiome associations

#### Overall procedure

CompDA adopts and modify recent statistical advances in fast and powerful high-dimensional conditional testing, i.e., the distilled conditional randomization testing method^27^, for providing valid inference for the conditional null hypothesis in **Definition 1**. Let *i* additionally denote sample indices. Briefly, for testing the conditional independence between *Y*_*i*_ and a given microbe *X*_*ij*_ in **Definition 1**, the procedure is as follows:

1. Perform predictive modeling of *Y*_*i*_ given 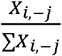, to obtain fitted values 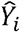. Also perform prediction modeling of (potentially transformed) *h*(*X*_*ij*_ ) given 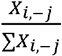 to obtain fitted values 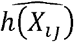 .*h* can be any transformation that is usually adopted for skewed microbiome relative abundances, such as the log transformation with pseudocounts or the arcsin-square-root transformation^7^. Let 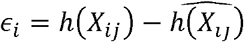 denote the residuals.
  a. The correlation statistic *T*(*Y*−*Ŷ*,∈ = | ∑_*i*_*∈*_*i*_(*Y*_*i*_−*Ŷ*) | thus quantifies the “strength” of association between *Y* and *X*_*j*_, while conditioning on 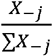. This is similar to e.g. regression coefficients obtained from partial least squares regression.
  b. *Y*_*i*_ given 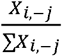 can be modeled with any prediction modeling approach including machine learning methods. In our implementation the default option is with (generalized) lasso regression, as implemented in the R package glmnet. The predictors are 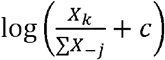 where *c* is a small positive value (pseudo count), set by default to half the minimum non-zero relative abundance.
  c. *h*(*X*_*ij*_ ) given 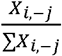 modeling requires careful consideration of microbiome data characteristics. This is discussed in more details in the next section.
2. Generate conditionally randomized versions of 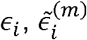 , for random iterations *m* =1,2,…,*M*, such that 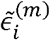 satisfies:
  a. Each 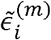 follows the same conditional distribution as 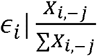
  b. 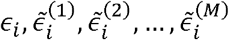 are mutually independent conditional on 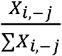 , and
  c. each 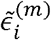 also satisfies that 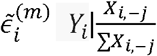 . Of the three conditions, 2.a. requires modeling of the distribution 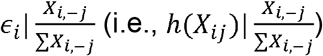 , whereas 2.b. and 2.c. are automatically satisfied if 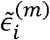 are generated as random variables independent of *Y*_*i*_.
3. With 2.a.-c. satisfied, the distribution of summary statistic *T* using the conditionally randomized samples,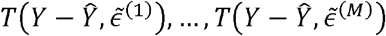 , forms a valid null distribution for the conditional independence hypothesis 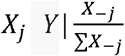 Thus a valid p-value can be calculated as 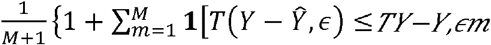 These individual p-values can additionally be synthesized with established FDR controlled procedures such as the Benjamini-Hochberg^35^ to prioritize true, health-related microbes while controlling false discoveries.

#### Modeling of *h*(*X*_*ij*_ )

The modeling for *h*(*X*_*j*_) given 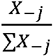 requires microbiome-specific consideration, as the relative abundance *X*_*j*_ is typically zero-inflated for microbiome data, which leads to a bimodal distribution for *h*(*X*_*j*_). To this end, we developed a component-wise prediction and residual conditional randomization procedure. Let 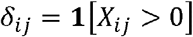 be the indicator that *X*_*ij*_ is positive. To obtain *∈*_*i*_ and 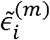 in the above procedure, the steps are:

1. To generate the residual *∈*_*i*_, we model separately either “mode” of *h*(*X*_*ij*_ ), corresponding to the zero and non-zero component *X*_*ij*_.
  a. Model *h*(*X*_*ij*_ ) given 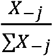 for when *X*_*ij*_.=0. This involves logistic lasso regression modeling of *δ*_*ij*_ given log-transformed 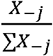 via glmnet, which yields a predicted value for the conditional probability 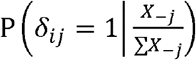 . We denote this by 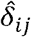.
  b. Model *h*(*X*_*ij*_ ) given 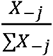 for when *X*_*ij*_ > 0. This involves linear lasso regression modeling of *h*(*X*_*ij*_ ) given log-transformed 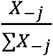 via glmnet, but only on samples where *X*_*ij*_ > 0. The fitted model nevertheless yields predicted values for all observations, which we denote by 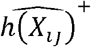.
  c. With the two components modeled, our predicted value for *h*(*X*_*ij*_ ) ,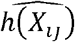 , is calculated as 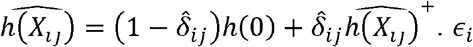 is correspondingly calculated as 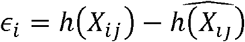
2. To generate the conditionally randomized residual 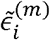 , for *m* ∈ {1,2,…,*M*} and each sample *i*, we also sample the zero and non-zero components separately:
  a. Sample zero/non-zero indicator 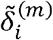 , where 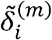 follows Bernoulli distribution with probability 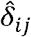.
  b. Sample the non-zero component 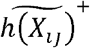 , by drawing a sample from the non-zero prediction modeling residuals in 1.b. That is, to sample from the residuals 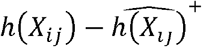 (obtained for only observations where *X*_*ij*_ > 0).
  c. With the two components sampled, our sampled value for *h*(*X*_*ij*_), 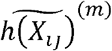 is calculated as 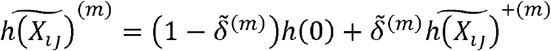 The conditional randomization residual 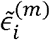 is correspondingly calculated as 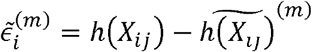 .

With this approach, we ensure that 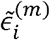 is not associated with *Y*_*ij*_ −*Ŷ*_*ij*_ but has the approximately same distribution as *∈* (both conditional on 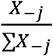 ). Thus 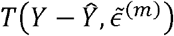 forms a valid null distribution for *T*(*Y*−*Ŷ* ,*∈*) corresponding to conditional independence between *Y* and *X*_*j*_.

### Accounting for host and environmental covariates

CompDA’s definition for conditional health-microbiome associations (**Definition 1**) and the accompanying testing procedures generalize easily to additionally account for host/environmental covariates. Let *Z* be the set of potentially confounding host and environmental factors (gender, diet, smoking history, etc.). **Definition 1** can be modified to the following, to additionally account for such factors:

#### **Definition 2**.

A microbe is conditionally not health-associated while additionally accounting for host and environmental factors, if its relative abundance is independent of the outcome while conditioning on the renormalized “balance” of the rest of the microbiome as well as the covariates *Z*. That is,

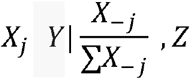

To test for this null, the above testing procedure can be extended by including *Z* in the conditional modeling of both *Y* and *h*(*X*_*j*_ ), in addition to 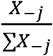.

### Simulation evaluation of method performance

We used real-world microbiome data to evaluate the performance of CompDA and compare against alternative methods. Specifically, we obtained metagenomic shotgun sequencing microbial samples of the human gut through the publicly available curatedMetagenomicData resource^30^. curatedMetagenomicData includes preprocessed microbial abundance observations from both diseased and healthy human subjects. For simulation studies, we filtered for healthy stool samples from westernized cohorts. This aggregated to a total of 8,000 samples and up to 300 microbial taxonomic features. Either binary or continuous health outcomes were simulated given these observations. Specifically, given our definition of health-microbiome conditional associations, a binary outcome *Y* is simulated via the following generalized linear model:

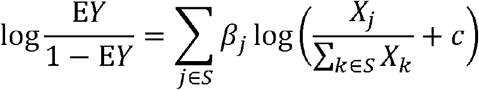

Here *S* is the set of true, health-related microbes. *c* is pseudo count to account for the inflated zeros in microbial abundance data, set to half the minimal non-zero relative abundance. Similarly, for a continuous outcome, the following model is adopted:

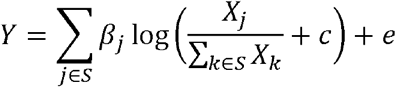

Where *e* is i.i.d. noise term with standard normal distribution. We note that for either simulation setting, the microbiome data *X* are directly obtained from real-world studies, thus rendering our simulation findings more generalizable to real-world applications.

We evaluate the full set of simulation parameters, by:

- Varying magnitude of health-microbiome association effects *β*_*j*_ For a prescribed effect size *l*, we sample *β*_*j*_ ∼ Uniformh(−2*l* ,2*l*) for *j ∈ S* That is, effects of true health related microbes are sampled from a uniform distribution.
- Varying sample size, i.e., the total number of observations in the data.
- Varying the complexity of the microbiome environment, i.e., the total number of microbes in each dataset.
- Varying signal density. Signal density is defined as the proportion of true health-related microbes compared to the total number of microbial features.

For each parameter specification, a total of 200 random replicates were generated (20 random subsets of real-world microbiome samples, multiplied by 10 random interactions of simulated health outcome *Y*). We synthesized performance of various methods and reported their false discovery rates and power for detecting true, health-related microbes for comparison.

We included the following methods for comparison:

- Traditional DA testing, as implemented by the popular MaAsLin2 method^7^. The default modeling choices (total sum scaling, log transformation) were used for analysis.
- Compositionality-adjusted DA testing (LinDA^17^). LinDA corrects for the bias caused by compositional effect in DA testing as estimated with the mode of regression coefficients across microbes. Still, it performs per-microbe marginal tests as DA, and thus does not consider microbe-microbe confounding associations. We adopted all default options in LinDA.
- CompDA (our approach).
- Lasso regression without inference, i.e., variable selection with non-zero coefficients from lasso regression. Specifically, we perform (generalized) linear regression with L1 regularization on *Y* with high dimensional covariates log (*X*_*j*_+*c*) Non-null microbes are selected as those with non-zero fitted regression coefficients. For model regularization, the optimal regularization parameter is selected with five-fold cross-validation.
- Lasso regression with inference. This is adopted from the microbiome covariates regression method from^21^. The authors proposed an additional debiasing procedure on lasso fits to enable inference of the regression coefficients, thus generating p-values. We obtained p-values via the matlab implementation provided by the original publication. As this only the binary *Y* implementation is provided, evaluation is limited to this case.
- Fixed-X knockoff procedure paired with screening^23^. This is available for continuous outcome and thus evaluated only for this case. We adopted the original R implementation of the authors. Control parameters were set as consistent with the original publication (40% of observations used for screening; screening set size set to 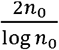 where *n*_0_ is the number screening observations). The fixed-X knockoff framework has restrictions regarding sample size (sample size must be no less than two times the number of features), and the authors aimed to address this limitation with their screening procedure. As such, we focused our evaluation for this method on varying the sample sizes (number of microbes at 200; number of samples at 50, 100, 200, and 400).

### Additional simulation studies

#### Misspecified health-microbiome association simulation with different machine learning models

For any modelling choice, CompDA provides statistically valid p-values for features selected by microbiome machine learning models. We selected to evaluate three specific models: (generalized) lasso regression (the default option), support vector machine (radial basis function kernel), random forest, and gradient boosted tree. The first model is implemented via the R package glmnet and the last three are implemented via the R package caret which provides wrappers around the R packages randomForest, kernlab, and xgboost, respectively.

We focus our evaluation for the binary *Y* case. Similar to the simulation scheme evaluated above, we simulated the outcome to be associated with log transformed, renormalized microbial abundances, but after taking the squares of the predictor values:

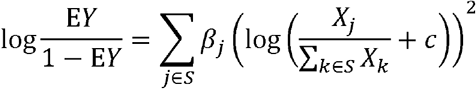

With this setup, our true data generation mechanism does not exactly agree with any of the underlying assumptions of the three machine learning models (strongly violates the assumptions of lasso and support vector machine, less so for random forest and gradient boosted tree). We then evaluate the performance of the three models and validate that they are controlled for false discoveries even for such misspecified scenarios.

#### Testing CompDA’s performance for adjusting for confounding variables

For simulating microbiome-health relations that are confounded by age, we obtained age group (*Z*) information from curatedMetagenomicData. These are matched with the microbiome profiles, thus the true associations between age and microbiome are naturally preserved in the simulation paradigm. We focus on the continuous *Y* scenario, and the outcome is simulated with:

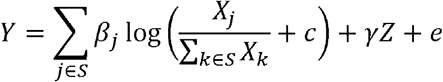

That is, the microbiome-health association is simulated same as before, but now the health outcome is additionally associated with the confounding age variable *Z · Z* is specifically defined as infancy versus others (i.e, age ≤ 2 or > 2), to reflect the strong difference between the infancy microbiome versus other stages of life.

We fixed the average microbiome effect size *β*_*j*_ ∼ Uniformh(−2,2) while varying the strength of the confounding age-health associations *γ*. The relative effect size of age compared to the microbiome on the outcome, as illustrated in **Figure 3B**, is defined as standard errorh ; (*γZ*)/standard error 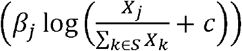 , i.e., the relative “contribution” to the variability in *Y* of age versus the microbiome. We evaluated the performance of CompDA analysis unadjusted for age versus age-adjusted CompDA analysis.

### Applications to real-world microbial signatures for colorectal cancer

We applied CompDA to identify microbial signatures of colorectal cancers (CRC) in a collection of colorectal microbiome studies^26^. Uniformly processed microbial relative abundance profiles are obtained again from curatedMetagnomicData^30^, including studies “ZellerG_2014”, “YuJ_2015”, “FengQ_2015”, “VogtmannE_2016”, “HanniganGD_2017”, “ThomasAM_2018a”, and “ThomasAM_2018b”. We applied CompDA to test for health-microbiome associations, including study-specific effects as potential confounders in modeling. Similarly, differential abundance testing was performed by regressing individual microbial abundances against both CRC status and study effects using the MaAsLin2 software with its default log transformation option^7^. Significant taxa were selected at an FDR threshold of 0.05 for both methods.

For examining the association of each microbe with CRC status while accounting for the rest of the microbe, we adopted the conditional association statistic adopted for CompDA testing, i.e., *T* (*Y*− *Ŷ*, *∈*)= | ∑_*i*_ *∈*_*i*_ (*Y*_*i*_ − *Ŷ*) | as reported in section “Conditional randomization testing of multivariable health-microbiome associations”. For examining the predictive power of *D. pneumosintes, F. nucleatum, P. stomatis, and S. salivarius*, we fitted regularized generalized linear prediction models using a) study indicator only as a baseline predictor, b) study + single microbe, c) study + all four microbes, and d) study + all microbes except for one. Model prediction performance (area under the ROC curve) was evaluated using nested five-fold cross validation. The difference in AUC between b) and a) indicates the individual predictive power of each microbe, while not accounting for the other microbes. The performance decrease from c) to d), however, indicates how important each microbe is, *given that the other microbes were included in the model*. It would thus distinguish microbes that are only spuriously related to CRC, as those that do not contribute to predictive power given that other microbes were included.

### Software and data availability

CompDA is implemented as an R package, available at https://github.com/syma-research/CompDA. The code to reproduce all analyses in this paper is available as a GitHub repository: https://github.com/syma-research/CompDA_paper.

## Acknowledgments

The computations in this paper were run in part on the FASRC Cannon cluster supported by the FAS Division of Science Research Computing Group at Harvard University.

