## Supplementary figures and images for "Compositional Differential Abundance Testing: Defining and Finding a New Type of Health-Microbiome Associations"

### Supplemental Figure 1

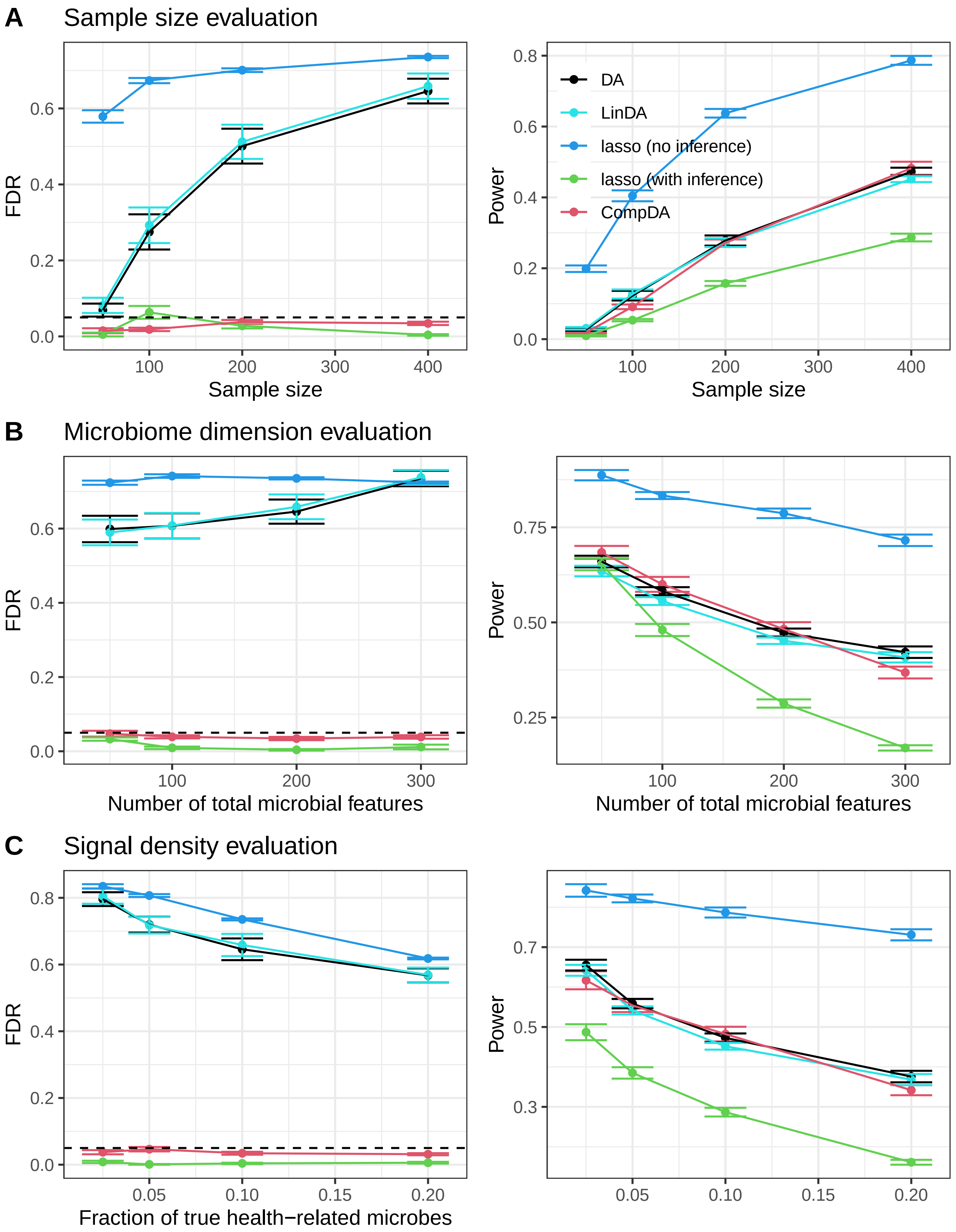

### Supplemental Figure 2

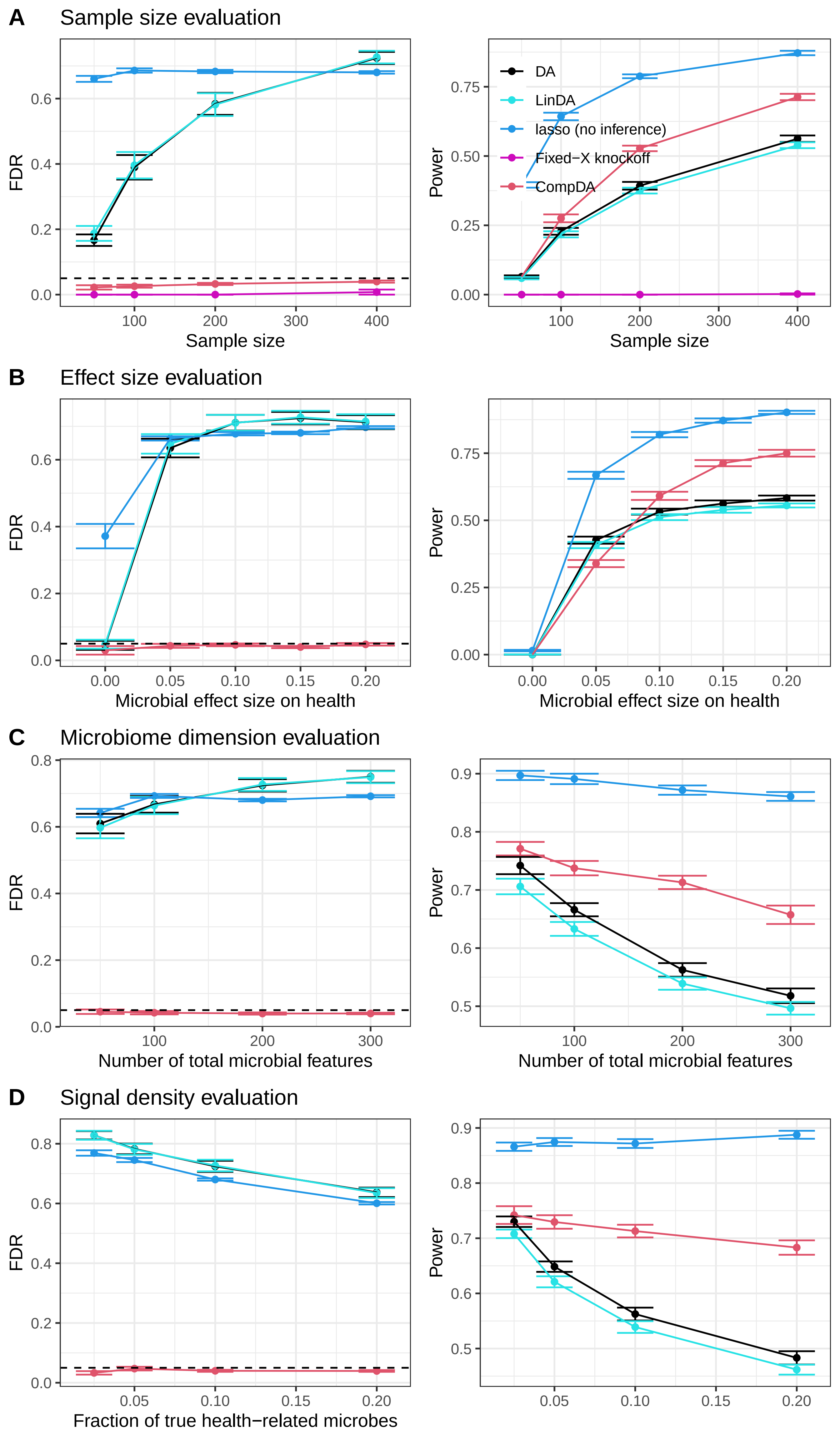

### Supplemental Figure 3

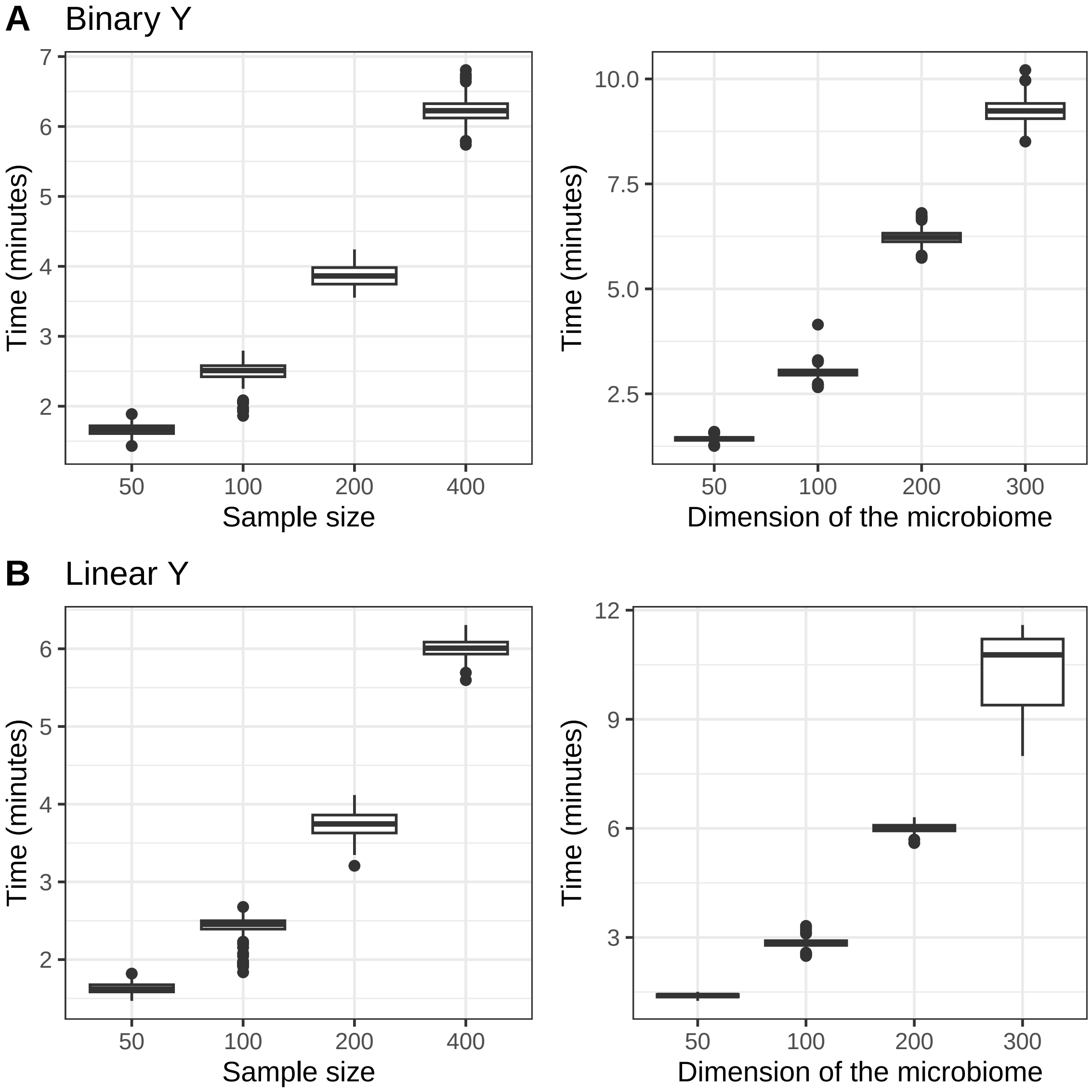
