## Supplemental Figure Captions for "Compositional Differential Abundance Testing: Defining and Finding a New Type of Health-Microbiome Associations"

**Supplemental Figure 1: Performance evaluation for CompDA and other methods for binary** $\boldsymbol{Y}$**.** In addition to varying the microbiome effect size on health (**Figure 2**), we also varied: **A**) the sample size, while fixing microbiome effect size at 0.4, number of features at 200, and signal density at 0.1; **B**) the microbiome complexity (number of microbes), while fixing the number of samples at 400, the microbiome effect size at 0.4, and signal density at 0.1; and **C**) the signal density (i.e, percentage of the microbes that are truly associated with health outcome), while fixing the number of samples at 400, the microbiome effect size at 0.4, and the number of microbes at 200. Across all simulation scenarios, CompDA successfully controlled the FDR while achieving the best power among methods that are able to do so.

**Supplemental Figure 2: Performance evaluation for CompDA and other methods for continuous** $\boldsymbol{Y}$**.** Similar to the binary $\boldsymbol{Y}$ evaluation, we examined CompDA’s performance varying one particular aspect of health-microbiome relationships at once while fixing other components constant. These include: **A**) the sample size, while fixing microbiome effect size at 0.15, number of features at 200, and signal density at 0.1; **B**) microbiome effect size, while fixing sample size at 400, number of features at 200, and signal density at 0.1; **C)** the microbiome complexity (number of microbes), while fixing the number of samples at 400, the microbiome effect size at 0.15, and signal density at 0.1; and **D**) the signal density, while fixing the number of samples at 400, the microbiome effect size at 0.15, and the number of microbes at 200. Again, across all simulation scenarios, CompDA successfully controlled the FDR while achieving the best power among methods that are able to do so.

**Supplemental Figure 3: Computational performance of CompDA.** We bench marked the computation cost of CompDA in both binary (panel **A**) and continuous (panel **B**) $Y$ settings across different sample sizes and number of microbes. Sample size was fixed at 400 while varying the number of microbes, and the number of microbe was fixed at 200 while varying sample sizes. Across all scenarios (which are likely to occur for real-world microbiome studies), CompDA can be completed with single core computing in under or around ten minutes. This can be improved in practice with parallelized computing, which we have implemented in the CompDA package. Benchmark was performed using Intel(R) Xeon(R) Gold 6134 CPU @ 3.20GHz.
